# Extensive phenotypic plasticity in a seasonal butterfly limits potential for evolutionary responses to environmental change

**DOI:** 10.1101/126177

**Authors:** Vicencio Oostra, Marjo Saastamoinen, Bas J. Zwaan, Christopher Wheat

**Author notes:** Shared senior authors (Bas J. Zwaan, Christopher Wheat).

## Abstract

Understanding how populations adapt to changing environments is a major goal in evolutionary biology and ecology, and particularly urgent for predicting resilience to climate change. Phenotypic plasticity, the ability to express multiple phenotypes from the same genome, is a widespread adaptation to short-term environmental fluctuations, but whether it facilitates adaptation to environmental change on evolutionary timescales, such as those under climate change, remains contentious. Here, we investigate plasticity and adaptive potential in an African savannah butterfly displaying extensive plasticity as adaptation to predictable dry-wet seasonality. We assess the transcriptional architecture of seasonal plasticity and find pervasive gene expression differences between the seasonal phenotypes, reflecting a genome-wide plasticity programme. Strikingly, intra-population genetic variation for this response is highly depleted, possibly reflecting strong purifying selection in its savannah habitat where an environmental cue (temperature) reliably predicts seasonal transitions and the cost of a mismatched phenotype is high. Under climate change the accuracy of such cues may deteriorate, rendering dominant reactions norm maladaptive. Therefore, depleted variation for plasticity as reported here may crucially limit evolutionary potential when conditions change, and seasonally plastic species may in fact be especially vulnerable to climate change.

## Introduction

Understanding how populations adapt to changing environments is of fundamental importance for assessing their evolutionary and ecological dynamics. This question has particular urgency for predicting population resilience to current climate change. Phenotypic plasticity, the expression of different phenotypes from the same genome in response to environmental change, is a widespread and important adaptation to habitats with fluctuating ecological opportunities. In seasonal habitats, ubiquitous in both tropical and temperate regions, phenotypic plasticity is particularly relevant, allowing organisms to maximize fitness as they track the predictable cycles of contrasting ecological conditions ^1^. These cycles place divergent selective pressures on an organism’s life history strategy, rewarding phenotype-environment matching via seasonally plastic strategies such as migration or diapause ^2^.

While the ecological relevance of plasticity is evident, its on-going impact on evolutionary dynamics is more contentious ^3-6^. Plasticity allows individuals to adjust to variable environments during their lifetime, but whether it helps populations to adapt to environmental change on evolutionary timescales remains to be resolved. Notably, it is unclear to what extent and under what circumstances phenotypic plasticity potentiates evolutionary adaptation to novel environments and thus facilitates population persistence under environmental change, or instead limits such adaptation. This question has particular urgency in the context of current rapid climate change, which is already having profound impacts on biodiversity and ecosystem functions ^7-9^ but where the integration of biological mechanisms into predictive models of resilience to climate change is hampered by a lack of empirical data, for example on physiology, phenology, or genetic variances ^10^.

One potential limit to adaptability in the face of environmental change that is particularly relevant in seasonal habitats is lack of genetic variation for plasticity ^9,11^. For seasonally plastic organisms it is crucial to match the expressed phenotype to what is needed to survive and / or reproduce in the prevailing seasonal environment. This requires predictability of seasonal transitions, and that the phenotypic response (reaction norm) is tuned to the reliability of an environmental cue as a predictor for the selective environment ^12,13^. If cue reliability is high, selection favours a reaction norm with high sensitivity to the relevant cue, i.e. a steep slope, and alternative reaction norms (based on unreliable environmental cues, or insufficiently sensitive to a reliable cue) will result in a maladaptive mismatch between phenotype and environment ^2,13,14^. Under stable long-term climatic conditions, this ongoing purifying selection acts to reduce standing genetic variation for plasticity from the population.

Under climate change, charactarised by increased environmental stochasticity ^15^, the reliability of many existing environmental cues as predictors for seasonal progression will likely diminish. As a consequence, the dominant reaction norm is more likely to produce mismatches between phenotype and environment, and this has been shown to produce maladaptive phenological shifts in many species ^16,17^. Under these conditions, selection will favour alternative reaction norms, for example with a different sensitivity to the environmental cue, or bet-hedging strategies ^18^. However, if past selection has depleted standing genetic variation for plasticity, the potential for evolutionary change will be limited, at least in the short term ^19^, reducing probability of population persistence under rapid climate change ^13,20^. Thus, rather than giving rise to adaptable generalists^21^, phenotypic plasticity in seasonal habitats may generally result in specialists with reduced long-term adaptive potential that are particularly vulnerable to climate change. Novel or stressful conditions, which under climate change are becoming more frequent ^15^, might aid adaptive potential, either by inducing phenotypic expression of genetic variance that is not visible under normal conditions ^22,23^, but see ^24^, or by disrupting normal patterns of plasticity and thereby producing novel phenotypes or combinations of phenotypes.

In order to better understand the role of plasticity in adaptation and resilience to rapid climate change, we use a textbook model of seasonal plasticity: the African savannah butterfly *Bicyclus anynana* ^25^. In the warm wet season, when food and reproductive opportunities are abundant, butterflies express a life history of fast growth and maximal reproduction. In contrast, during the cool dry season, when adult food is limited and larval food completely absent, the life history phenotype is focused on inactivity and postponed reproduction ^26-28^. The seasonal phenotypes also include alternative wing and behavioural patterns, and are reliably induced via temperature cues in laboratory populations ^25^. As they are produced by deploying the shared genome differently between the seasons, we use a transcriptomic approach and characterise the transcriptional architecture of the seasonal dimorphism ^29^.

Using this species, we assess the potential for evolutionary change in plasticity by characterising the transcriptional architecture of the seasonal phenotypes, quantifying intrapopulation genetic variation for plasticity, and determining the role of stress in adaptability. We uncover high amounts of seasonal plasticity in the transcriptome, reflecting a genome-wide plasticity programme, consistent with the broad suite of traits involved in the seasonal adaptation. This programme is partly systemic, and partly tissue-specific, reflecting tissue-specific life history trade-offs across environments. Strikingly, intra-population genetic variation for this plastic response is highly depleted, with low levels of gene-by-environment interaction and highly similar seasonal responses across different full-sib families. This depletion possibly reflects strong purifying selection in its savannah habitat where an environmental cue (temperature) reliably predicts seasonal transitions and the cost of a mismatched phenotype is high ^30^. Stressful conditions during development subtly decrease the transcriptional divergence between the seasons, but this effect is very modest, and, crucially, not accompanied by increased genetic variation for plasticity. Our results illustrate that lack of genetic variation for plasticity may crucially limit evolutionary potential and population persistence under environmental change, suggesting that seasonally plastic species adapted to predictable environments may in fact be especially vulnerable to climate change.

## Materials and Methods

### Study organism and experimental design

We used a captive laboratory population of B. *anynana*, reared under standardised outbred conditions^25,31^. In order to assess the effect of genetic background, seasonal environment, food stress and their interactions on the transcriptome, we employed a full factorial split brood experimental design with seven full sib families, two rearing temperatures (19C corresponding to dry season conditions and 27C corresponding to wet season conditions), and two developmental food stress treatments (*ad libitum* and food stress). Within this same experiment we have collected phenotypic data on additional individuals and families, which we published previously ^32^ including additional details on the experimental setup. Larvae completed their full development in one of two temperature conditions representing the seasonal environments, while the developmental food stress treatments lasted two or three days during the fifth (last) larval instar ^32,33^, with control larvae receiving normal food (fresh maize leaves) and stressed larvae receiving no food, only agar to avoid dehydration. One day after eclosion we sampled females for RNA isolation (snap-freezing in liquid N_2_), separating thorax and abdomen for separate sequencing.

### RNA isolation, sequencing and data pre-processing

We isolated RNA using TRIzol (Invitrogen) followed by the RNeasy Mini Kit (Qiagen) and a DNA digestion using the RNase-Free DNase enzyme (Qiagen). We sent total RNA for 144 samples (thorax and abdomen of 72 females) to BGI (People’s Republic of China) for library preparation and sequencing (paired-end, 2x 100 bp, average insert size 350 bp, Illumina HiSeq 2000). We obtained on average 15.4 × 10^6^ raw PE 100 bp reads per sample (95% CI: 13 – 16 × 10^6^), totalling 2.2 10^9^ reads across 144 samples. We assessed quality using FastQC v. 0.10.1 (http://www.bioinformatics.babraham.ac.uk/projects/fastqc/) and trimmed reads using bbduk2 in bbmap v. 0.34.94 (http://bbmap.sourceforge.net/), trimming on average 6.9% of reads per sample (95% CI: 4.5 – 10.7 %). See Supplementary Table S1 for an overview of sequencing results for all libraries.

### De novo transcriptome assembly and annotation

Transcriptome assembly used Trinity v. 20140717 ^34^, combining reads from all libraries, yielding 496,087 contigs with an N50 of 519 and an N10 of 2,395. Because *de novo* transcriptome assemblies produce a large number of false or chimeric contigs, resulting from transcriptional noise and misassemblies, we enriched our assembly for biologically meaningful transcripts using the EvidentialGene pipeline (http://arthropods.eugenes.org/EvidentialGene/trassembly.html). This yielded a filtered high quality transcriptome of 35,747 contigs with an N50 of 1,839 and N10 of 4,561. To annotate it we performed a blastp search of the 35,747 predicted proteins against the UniRef90 database ^35,36^. After filtering (evalue < 0.00001 and bitscore > 50), this yielded valid UniRef protein names for 14,681 transcripts. We then used Argot2 to obtain Gene Ontology (GO) terms for our transcriptome ^37^, yielding annotations for 15,991 transcripts, with on average 5.2 GO terms per annotated transcript.

### Mapping and transcript abundance estimation

We mapped trimmed reads against our filtered transcriptome using Bowtie2 v. 2.2.3 ^38^ allowing one mismatch between seed and reference, with an average of 83% reads mapped (95% CI: 72 – 93%; Supplementary Table S1). We quantified raw read counts per transcript per library using SAMtools idxstats ^39^. To further reduce transcriptional noise prior to differential expression analysis, we removed genes with low or very limited expression, processing the 72 thorax and 72 abdomen libraries separately. We removed genes that were expressed in less than 3 samples as well as genes with average expression < 0.25 counts per million (CPM). After filtering we retained 15,049 and 12,567 genes for abdomen and thorax, respectively, accounting for more than 99.9% of all read counts.

### Differential expression analysis

We performed differential expression analyses in edgeR v. 3.10.0 ^40^ to test the main effects of seasonal temperature, food treatment, family, as well as the three two-way interactions and the three-way interaction between the main effects on expression of all expressed genes in the transcriptome (Supplementary Table S2). As input data for edgeR we used untransformed, raw count data (after filtering to remove genes with low expression), and we performed all analyses separately for thorax and abdomen samples. For each factor of interest, edgeR analyses yielded a fold change between the conditions, a likelihood ratio for the effect of that factor, and a corresponding p value. We corrected these p values for multiple comparisons using Benjamini and Hochberg’s False Discovery Rate FDR ^41^, accepting an FDR of 0.05. The sign of fold change values for the effect of season is with reference to the wet season, i.e. genes with positive and negative fold change values being wet and dry season-biased, respectively.

### Gene Ontology analyses

We performed Gene Ontology (GO) Gene Set Enrichment (GSE) analyses in Babelomics v. 5 ^42^, using the FatiScan module ^43^. As input we used lists of genes (and their fold change) that were significantly season-biased in thorax, abdomen, or both body parts, and the Argot2 GO annotation file (after removing GO terms with with < 4 or > 500 genes, and terms with total score < 200). The GSE analyses yield a log odds ratio for enrichment associated with each GO term, with the same sign as fold change values in expression: positive for GO terms enriched among wet season-biased genes and negative for GO terms enriched among dry season-biased genes. The associated p values are corrected for multiple comparisons ^41^. We report only GO terms with FDR < 0.1 and more than 3 genes associated with it. To summarise and visualise long lists of GO terms we used REVIGO ^44^ with the following parameters: default similarity (0.7), default semantic similarity measure (SimRel), *Drosophila melanogaster* database, including the log odd ratios from the GSE analyses (with higher absolute values is better). We visualised the output in scatterplots.

### Cluster analyses, Principal Components Analyses (PCA), reaction norms, and cross-environment genetic correlations

For downstream analyses we normalised expression data in edgeR v. 3.10.0 ^40^ using trimmed methods of means (TMM), transformed to counts per million (CPM) and subsequently log_10_ transformed (for cluster analysis and PCA) or Z transformed (for reaction norm analyses and cross-environment genetic correlations). Cluster analyses were performed by constructing a neighbour joining tree from the Euclidian distance matrix computed from the normalised expression data using the R package “ape” ^45^. PCA were calculated by single value decomposition using the R function prcomp. To assess the association between Principal Components (PCs) and experimental factors, two-way ANOVAs were performed on each PC separately, with seasonal environment, family, and their interaction as fixed factors. We computed family-specific reaction norms for each gene by fitting, for each family separately, the normalised expression data for that gene in a general linear model with season as the sole predictor. This yielded, for each family separately, the intercept and slope of this model, which was then used to calculate coefficient of variance across the seven full-sib families for both the intercept and slope. Differences in the transcriptome-wide distribution of these coefficients of variance were tested with Wilcoxon signed rank tests (using the function wilcoxsign_test from R package “coin”). We calculated cross-environment genetic correlations in normalised expression for each gene by averaging gene expression per family within each seasonal environment (N = 4-6) and calculating Pearson’s correlation coefficient using the expression of each of seven families in the wet season and in the dry season. Differences in log_2_ fold change value between groups of individuals and groups of genes were tested with Wilcoxon signed rank tests. Log_2_ fold changes were calculated in edgeR, and those involving family (as main effect or in interaction with another factor) were calculated using all six mutually orthogonal contrasts between the seven families, and averaged across contrasts. Using maximum log_2_ fold change across contrasts rather than average yielded similar results.

All statistical analyses were performed in R v. 3.0.2 ^46^.

## Results

### Seasonal plasticity across the transcriptome

In order to understand how the shared genome is deployed differently across the seasonal environments to produce the distinct phenotypic morphs, we analysed the transcriptional architecture of plasticity by comparing gene expression between wet and dry season individuals. Three separate analyses revealed large fractions of the abdomen and thorax transcriptome to be involved in seasonal plasticity. Differential expression analyses revealed that in abdomen and thorax, 46 and 47% of genes showed significant season-biased expression, respectively (FDR < 0.05; Figure S1). Principal Components Analysis (PCA) of the whole-transcriptome expression profiles found PC1, accounting for 15-16% of total variance, significantly separating individuals from wet and dry season environments (two-way ANOVA1,55 F > 38, FDR < 10^−6^; Figure 1; Supplementary Figures S2, S3). Hierarchical clustering of gene expression confirmed these patterns, with individuals clustering strongly by season (Supplementary Figure S4).

**Figure 1.**
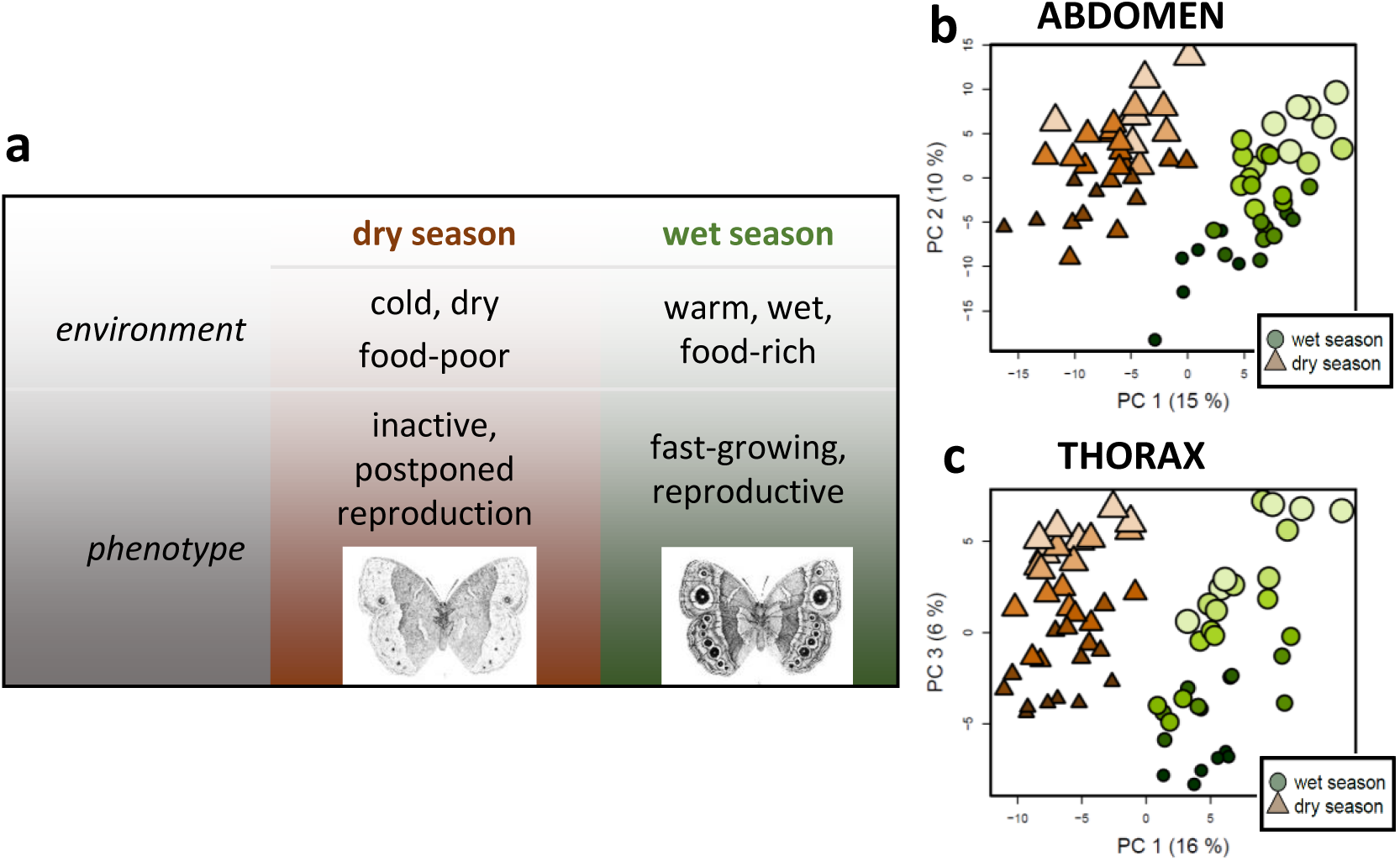
Pervasive seasonal plasticity and intra-population genetic variation across the transcriptome. **(a)** *Bicyclus anynana* butterflies of the dry (brown) and wet (green) season differ in a suite of behavioural, life history, and morphological traits, depending on the seasonal environment in which they developed.**(b,c)** Principal Components Analysis (PCA) of whole-transcriptome expression profiles for abdomen **(b)** and thorax **(c)** significantly separates individuals reared in wet (green circles) or dry (orange-brown triangles) season conditions along the first Principal Component (accounting for 15-16% of variance). For abdomen, the second PC (10% of variance) significantly separates individuals from different full-sib families (indicated by different colour shades and size of symbols), whereas for thorax the third PC (6 % of variance) separate families. Families are also separated by several other PCs, together accounting for 56 and 27% of variance in abdomen and thorax (see Supplementary Figure S2 and S3 for details).

Examining the seasonal plasticity programme in more detail revealed tissue-specific and systemic components, both at the level of individual genes, functional processes, and the whole transcriptome (Supplementary Results). A total of 2,093 genes showed the same response to the seasonal environment in both body parts, representing 14 and 17% of the abdomen and thorax transcriptome, respectively, and enriched for 89 functional GO terms. In addition to these systemic plasticity genes, 32 and 30% of genes, enriched for 155 and 189 GO terms, showed tissue-specific seasonal bias in abdomen and thorax transcriptomes (Figure 2; Supplementary Table S3; Supplementary Figure S5). Of the overrepresented GO terms identified separately in abdomen or thorax, 37 were shared among the tissues, representing 865 additional season-biased genes. This reflects an additional systemic signature of seasonal plasticity only apparent at the level of functional processes, not individual genes (Supplementary Table S3). Finally, seasonal expression bias in abdomen is positively associated with season bias in thorax (Figure 2B), with the fold change in one body part explaining 15% of that in the other body part (ρ_pearson_ = 0.38, t_12307_ = 46.032, p < 10^−15^; ρ _spearman_ = 0.27, t_12307_ = 30.7006, p < 10^−15^), indicating an additional quantitative systemic signature of seasonal plasticity across thousands of genes in the transcriptome. Thus, within and across tissues, the seasonal environment is a major determinant of transcriptional variation, representing a genome-wide seasonal plasticity programme (Supplementary Results), consistent with the broad suite of phenotypic traits involved in the seasonal adaptation ^28,32^.

**Figure 2.**
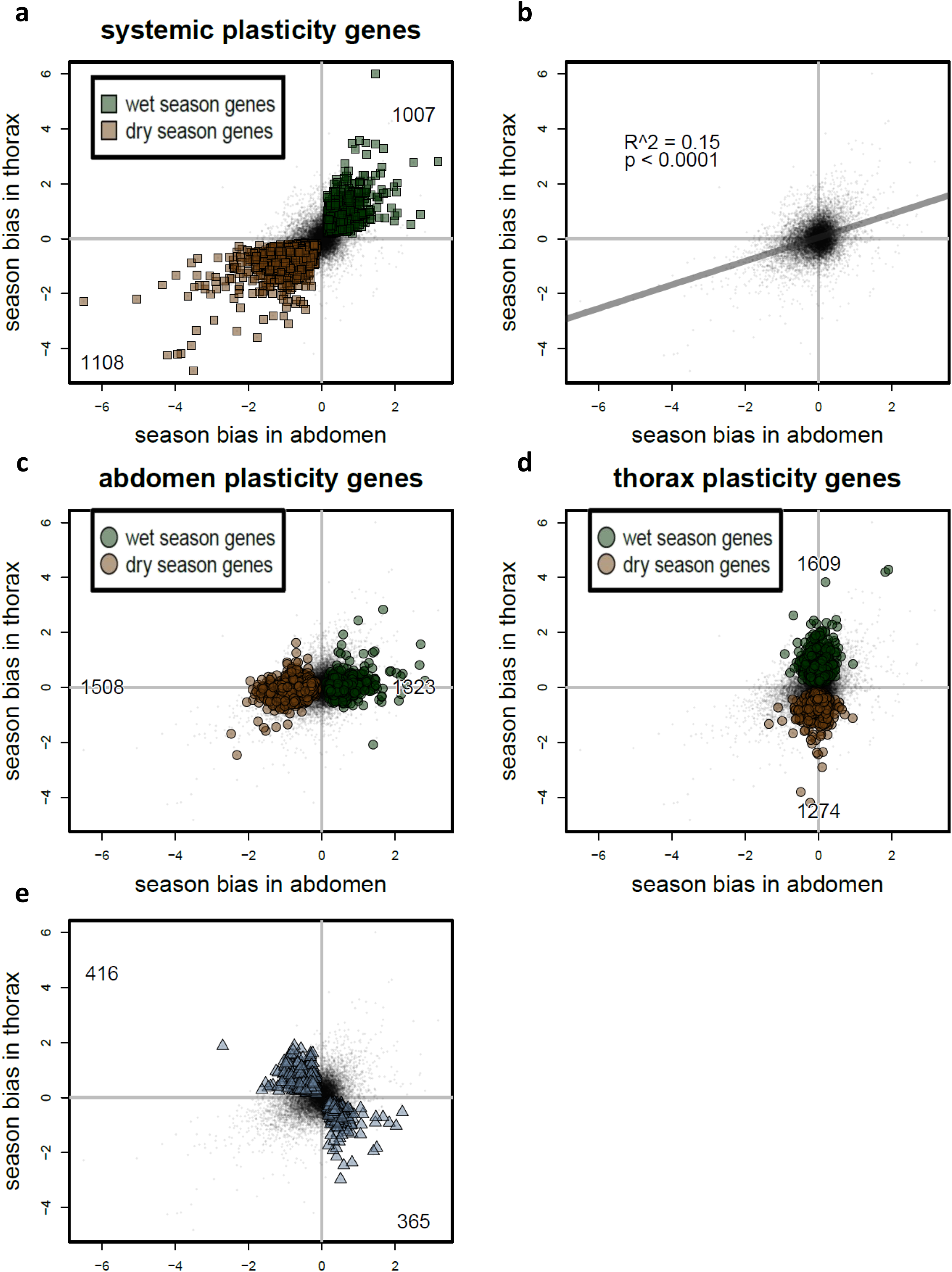
Systemic and tissue-specific components of the plasticity programme. For each gene expressed in both body parts, seasonal expression bias (log_2_ Fold Change of expression in wet season compared to dry season) in thorax (vertical axis) is plotted against expression bias in abdomen (horizontal axis), with positive Fold Change values indicating higher expression in wet season, and negative values indicating higher expression in dry season. Each gene is represented by a black dot, and genes significantly differentially expressed (FDR < 0.05) between wet and dry season are coloured in green and brown for wet and dry season-biased genes, respectively (**a, c,** and **d**), or in blue (**b**). Numbers in each quadrant indicate numbers of significant wet or dry season-biased genes. **(a)** The systemic plasticity programme is represented by 2,093 genes that show the same response to the seasonal environment in both body parts, representing 14 and 17% of the abdomen and thorax transcriptome, respectively, and enriched for 89 functional GO terms. **(b)** In addition to the significantly season-biased genes, body parts also show quantitative, transcriptome-wide overlap in seasonal plasticity. Seasonal expression bias across all expressed genes is significantly correlated between the body parts (ρ_pearson_ = 0.38, t_12307_ = 46.032, p < 10^−15^; ρ_spearman_ = 0.27, t_12307_ = 30.7006, p < 10^−15^), with expression bias in one body part explaining 15% of variation in expression bias in the other body part. **(c, d)** The abdomen- and thorax-specific plasticity programmes are represented by 2,831 and 2,883 genes, respectively, representing 32 and 30% of all genes and enriched for 155 and 189 GO terms. **(e)** Expression for 718 genes shows opposite patterns of season bias between the two body parts, i.e. expression is wet season-biased in one body part and dry season-biased in the other body part. These genes are enriched for 37 GO terms. For functional analyses of biological processes underlying the systemic and tissue-specific plasticity programme, see Supplementary Table S3, and Supplementary Figure S5.

### Depleted genetic variation for plasticity

The extent to which the transcriptional plasticity programme can evolve, for example in response to climate change, critically depends on standing genetic variation for plasticity in gene expression. We found, based on five different analyses, that this variation is highly depleted within the population, contrasting starkly with the high levels of standing variation for average expression.

First, expression of only 1% of genes (160 and 146 genes in abdomens and thoraces, respectively), was significantly affected (FDR < 0.05) by the interaction between seasonal environment and family, i.e. genotype-by-environment interaction (GxE). In contrast, genetic background (i.e. family) significantly affected average expression for 66% and 42% of genes in abdomen and thorax, respectively (Figure 3A, B; Supplementary Figure S1; Supplementary Table S4).

**Figure 3.**
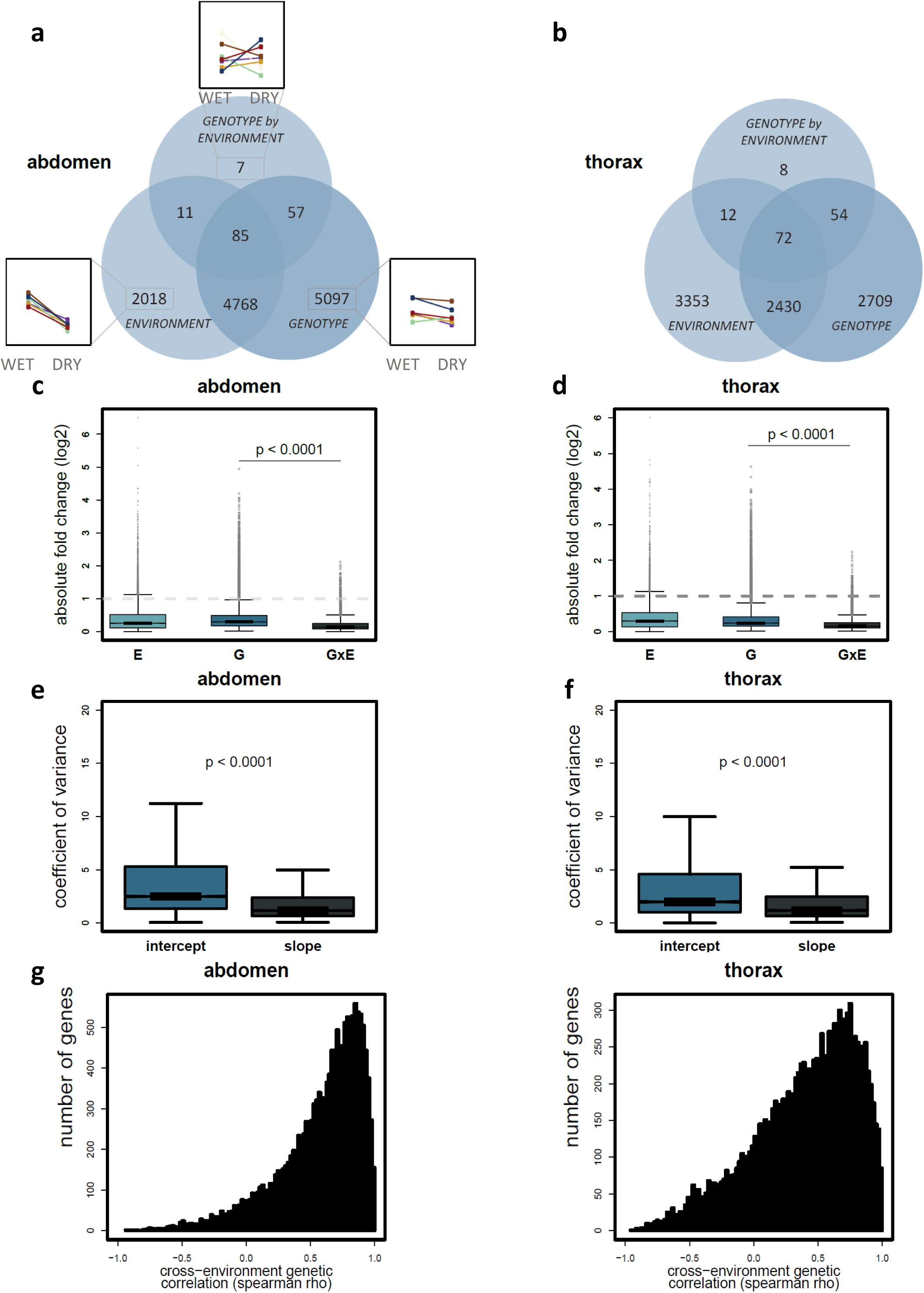
Large-scale depletion of genetic variation for seasonal plasticity across the transcriptome. **(a, b)** An order of magnitude more genes show significant differential expression due to seasonal environment and genetic background than due to the interaction between environment and genetic background for abdomen **(a)** and thorax **(b)**. Within each Venn diagram, numbers of differentially expressed genes (FDR < 0.05) are indicated for seasonal environment (left), genetic background (right), and their interaction (top), as well as overlap in responses among genes. Small insets in **(a)** illustrate expression patterns for each gene group, showing average normalised expression in wet and dry season for a random gene in each group, with reaction norms for different full-sib families represented by different colours. **(c, d)**. Across all genes in the abdomen **(c)** and thorax **(d)** transcriptome, absolute expression changes (log_2_ fold change) due to the interaction between environment and genetic background (i.e. transcriptional plasticity or GxE; right boxplot) are 43 - 51% smaller than those due to genetic background (i.e. genetic variation for average expression or G; middle boxplot). 120 and 88 transcripts had an absolute fold change higher than two for the gene-by-environment interaction, compared to 1,054 and 766 transcripts (in abdomen and thorax, respectively) above that threshold for mean inter-family expression differences. **(e, f)** Families are more similar in how expression responds to the environment than in average expression levels, with reaction norms for gene expression showing 57 and 32% lower across-family variance in slope than in intercept for abdomen **(e)** and thorax **(f)**, respectively. Boxplots show transcriptome-wide distributions of across-family variation (coefficient of variance) in intercept (left, blue) and slope (right, black). **(g, h)**. Across-environment genetic correlations in expression are positive and high for both abdomen **(g)** and thorax **(h)**. The histogram shows the distribution across all genes of Pearson’s correlation coefficients for the correlation between average per-family expression across the two seasonal environments.

Second, independent of statistical significance testing, the absolute expression changes between families in transcriptional plasticity were 43-51% smaller than those between families in average expression (Figure 3C, D; Wilcoxon signed rank tests Z < −133, P < 10^−16^). 120 and 88 transcripts (in abdomen and thorax, respectively) had an absolute fold change higher than two for the gene-by-environment interaction, compared to 1,054 and 766 transcripts above that threshold for inter-family expression differences.

Third, PCA confirmed that genetic variation for seasonal plasticity represents a tiny fraction of total transcriptional variance, contrasting with overall genetic variation being a large driver of average gene expression. In thorax, none of the first 13 Principal Components (together accounting for 62% of total variance) associated with the interaction between seasonal environment and genetic background (two-way Anovas, FDR > 0.49, F < 2.0), while in abdomen only PC 13 (accounting for 1.5% of total variance) was significantly affected by the interaction between seasonal environment and genetic background (FDR = 0.03, F = 3.9). In contrast, 7 and 9 PCs, accounting for 56 and 27% of variance in abdomen and thorax, respectively, significantly associated with the genetic background (two-way Anovas; FDR = 0.01 – 10^−24^; F = 3.3 – 85.1; Supplementary Figure S2, S3).

Fourth, analysis of reaction norms for gene expression revealed that families are much more similar in how expression responds to the environment than in their average expression levels. We computed family-specific seasonal reaction norms, and comparing across-family variance in slope (plasticity) and intercept (average expression) found that variance in slope was 57 and 32% lower than variance in intercept for abdomen and thorax, respectively (Figure 3E-F, F; Z > 36, p < 10^−16^).

Finally, we observed high positive genetic correlations in gene expression across seasonal environments (Figure 3G, H), with median Spearman Rank correlations of + 0.67 and + 0.47 for abdomen and thorax, respectively. This indicates that evolutionary changes in expression will be hampered unless they occur in the same direction in each season, and thus that potential for evolutionary change in plasticity is limited.

Together, these analyses provide evidence for a substantial depletion of intra-population genetic variation for plasticity, with low levels of gene-by-environment interaction and different full-sib families having highly similar seasonal responses. This contrasts sharply with the high levels of genetic variation for expression averaged across the seasons, where families vary extensively in average expression patterns. As a consequence, there is ample opportunity for evolutionary change in average expression in both seasons simultaneously, but potential for change in plasticity is severely limited.

### Developmental food stress and adaptability

We assessed whether stressful conditions might aid adaptive potential, either by inducing phenotypic expression of genetic variance that is not visible under normal conditions, or by disrupting normal patterns of plasticity. We exposed developing larvae (of both seasonal forms) to a period of food stress, and observed only minor effects on seasonal plasticity (Supplementary Results; Supplementary Table S5). Expression of less than ten genes showed a significant interaction between seasonal plasticity and food stress, indicating that for virtually all genes the seasonal response was not different under stressed compared to *ad libitum* conditions (Supplementary Figure S1). Examining the effect of food stress in more detail for specific gene repertoires revealed subtle stress-induced shifts in normal seasonal expression patterns. In particular, stress pushed the typical dry season morph towards a slightly more wet season-like transcriptional profile, partly driven by a terminal reproductive investment in the abdomen, with wet season genes showing increased expression upon stress (Supplementary Results; Supplementary Figure S6). However, this limited reduction in transcriptional divergence between the seasonal forms was not accompanied by increased genetic variation for plasticity. The three-way interaction between genetic background, food stress treatment and seasonal environment only affected a small number of genes, indicating very limited effect of stress on genetic variation for plasticity and thus potential for stress to release genetic variation in plasticity (Supplementary Figure S1). Thus, we found no evidence that the novel environment represented by food stress releases hidden genetic variation and aids adaptablity.

## Discussion

Understanding the factors affecting evolutionary adaptation and population persistence in novel environments is a key goal in evolutionary biology, and an urgent requirement to predict biotic responses to climate change. The role plasticity plays in facilitating or constraining adaptability has been discussed extensively in the literature (e.g. ^3-6^). On the one hand, evolutionary adaptation in novel environments may be constrained due to plasticity reducing the strength of selection by buffering environmental variation and thus shielding low fitness variants from selection (e.g. ^8^). On the other hand, plasticity is hypothesised to facilitate evolutionary adaptation in novel environments as plastic populations express a wider range of phenotypes than non-plastic populations, and new phenotypic optima may be more easily reached if they can be directly induced by the environment rather than produced from new genetic variants. In either perspective, plasticity is assumed to be a generalist strategy (e.g. ^21,47^) while this is often not the case. In particular, seasonal environments usually vary predictably, selecting for plastic strategies that specialise in precisely tracking seasonal progression by tuning to the right environmental cues such as day length or temperature.

Here, we report on adaptive potential and seasonal plasticity in the tropical butterfly *B. anynana*, which inhabits a stable savannah environment where seasonal transitions are highly predictable. Although this species shows pervasive seasonal plasticity across its transcriptome, standing genetic variation for plasticity in the population is highly reduced, likely the result of past purifying selection for a single optimal reaction norm that precisely captures the habitat’s reliable seasonality pattern. If long-term climatic stability ends, reducing predictability of seasonal transitions, the current optimal reaction norm may become maladaptive and alternative reaction norms likely become more favourable. Given the lack of genetic variation for plasticity, the potential for an adaptive shift towards a new environmental response will be severely limited, with potentially serious negative consequences for population persistence under rapid climate change.

We observed pervasive seasonal plasticity across the transcriptome, with almost half of all expressed genes showing season-biased expression. This was partly driven by a systemic plasticity programme, and partly by tissue-specific regulation of gene expression that reflects tissue-specific life history trade-offs across environments. The involvement of a large portion of the transcriptome in seasonal plasticity is not surprising, given that the seasonal adaptation in *B. anynana* represents a broad behavioural and life history syndrome. This syndrome encompasses a suite of phenotypic traits, including growth rate, size at maturity, hormone physiology, metabolic rate, fat metabolism, reproductive strategy, and starvation resistance ^26-28,32,48-51^, consistent with the enriched functional processes among the season-biased genes.Remarkably, a separate study found the head transcriptome to be much less differentiated between the seasonal forms than our results for abdomen and thorax ^52^, possibly indicating that behavioural differences are less pronounced than physiological phenotypes, at least early in adult life. In other phenotypically plastic animals, the extent of transcriptional differentiation between alternative morphs varies widely, likely reflecting different levels of physiological or morphological specialisation into alternative phenotypes (e.g. ^53-59^).

Although seasonal plasticity is pervasive across the transcriptome, there is very little intra-population genetic variation for it, contrasting markedly with the extensive average variation for gene expression. The families differ significantly in average expression for approximately half of all transcripts, with high inter-family variance in reaction norm elevation. These high levels of variation for average gene expression have also been observed in other animal populations (e.g. in fruitflies^60^; butterflies^61^; fish^62^; baboons^63^; humans^64^; and other species in the wild:^65^). In marked contrast, the families display the same transcriptional response to the seasonal environment for virtually all transcripts, and inter-family variance in reaction norm slope is significantly reduced. The observed lack of variation for plasticity is unlikely due to inbreeding, as that would also depress overall expression variation. Another potential confounder, relaxed purifying selection due to laboratory breeding conditions, is also unlikely to play an important role. Relaxed selection on plasticity would likely result in increased rather than decreased genetic variation for plasticity ^66^ and potentially reduced plasticity overall cf. ^67^. Importantly, our results on transcriptional plasticity support earlier work on phenotypic trait plasticity and evolution on this same population. Many seasonally plastic traits, such as growth rate, metabolic rate, starvation resistance and wing pattern also show significant intra-population genetic variation ^33,48,68,69^, and artificial selection was highly successful in shifting mean trait expression, i.e. the reaction norm elevation (e.g. ^68^). In contrast, reaction norm slopes showed high across-environment genetic correlations and very limited genetic variation, precluding evolutionary responses to artificial selection targeting reaction norm slope ^70,71^. As the latter selection experiments were only performed for wing pattern, it remains to be tested whether life history traits are similarly constrained in potential for change in plasticity, but our transcriptional data strongly suggest that this indeed is the case.

Although food stress during development increases resilience to stress later in life in this species ^33,72^, developmental food stress had only a marginal effect on the transcriptome, and it did not increase additional genetic variation. While it slightly disrupted the plasticity programme, pushing the dry season form towards a slightly more wet season-like transcriptional profile, this effect was quite minor and was not accompanied by increased levels of genetic variation. The developmental food stress treatment lasted only 2-3 days, which may not seem severe, but earlier experiments show that an additional day of food stress increases mortality dramatically ^33^. Thus, unlike in some other studies ^22,23^, (but see ^24^), in our study stressful conditions did not release genetic variation and therefore do not aid adaptability under environmental change.

*Bicyclus anynana* inhabits a strongly seasonal savannah environment, with vastly differing food availability and reproductive opportunities between the dry and wet seasons, making it crucial to produce the right phenotype at the right moment. In Nkhata Bay in Malawi, where the laboratory population originates, seasonality is stable over the long term, with highly predictable transitions between the seasons ^27^. Available climate data for 1901-2009^73^ reveals a strong positive correlation between monthly-averaged precipitation and minimum temperature in the previous month (ρ_pearson_ = + 0.85; Supplementary Figure S7). This puts a premium on a single reaction norm optimally tuned to the specific temperature cue predicting seasonal transitions. Alternative reaction norms would produce either the wrong phenotype, or the right phenotype at the wrong time of the year, leading to a maladaptive mismatch between seasonal environment and expressed phenotype ^30^. Given the vastly contrasting selective pressures across the seasons, there is likely strong purifying selection removing such mismatch-producing reaction norms from the population, resulting in the observed depletion of genetic variation for plasticity without affecting overall levels of genetic variation for expression.

Our results indicate that the potential for evolutionary change in plasticity in this population is constrained by the current lack of genetic variation for plasticity, at least in the short term, and any evolutionary response in plasticity will thus depend upon novel mutations affecting reaction norms. This limited adaptability will only surface if selective pressures change in such a way that the currently dominant reaction norm no longer has the highest fitness, and instead produces a mismatch between expressed phenotype and the new selective environment. Current climate change represents exactly such a scenario, as not only temperature means are increasing rapidly, but also variance in temperature and precipitation ^15^. This will likely entail a decrease in the presently high correlation between temperature and rainfall and thus a deterioration in the reliability of existing environmental cues for seasonal progression, increasing selective pressures for alternative reaction norms.

In many seasonal habitats species depend on environmental cues such as temperature or day length to prepare for seasonal transitions ^47^. A modeling study shows that decreased environmental predictability increases extinction risk for plastic species ^20^, and indeed climate change, by affecting environmental cues as well as seasonal timing, is already increasing mismatches between phenotype and seasonal environment ^16,17,19,74^ but see ^75^. Even where species respond to climate change by shifts in seasonal timing, the adjustment in phenology is often too subtle or too extreme, leading to frequent mismatches in phenotype with seasonal progression ^17^. Although microevolutionary changes in such responses are crucial for population resilience to climate change, there is very little empirical data on the extent of genetic variation for plastic responses, limiting our understanding of adaptive potential of populations facing climate change ^8,9,19,76^. Our study illustrates that specialised seasonal plasticity may result in reduced adaptability in the face of environmental change via depletion of genetic variation for seasonal reaction norms. Given the ubiquity of seasonal habitats across tropical and temperate areas this likely applies to many species, and thus represents an underappreciated limit to biotic climate change resilience.

## Data availability

RNA sequencing reads are available at NCBI Sequence Read Archive, BioProject ID PRJNA376691, and the transcriptome assembly as well as Supplementary Table S3 are available at Figshare (doi: 10.6084/m9.figshare.4834031).

## Acknowledgements

This work was generously supported by the European Commission under FP7 (IDEAL FP7/2007-2011/259679 to B.J.Z. and V.O.) and Marie Sklodowska-Curie Fellowships (660172 to V.O.), by the European Research Council (Independent Starting grant META-STRESS; 637412 to M.S.), by the Academy of Finland (Decision numbers 273098 and 265641 to M.S. incl. Centre of Excellence funding to VO), by the Knut and Alice Wallenberg Foundation (KAW 2012.0058 to C.W.) and by the Swedish Research Council (VR-2012-4001 to C.W.). We thank Kees Koops, Mariël Lavrijsen,David Hallesleben and Erik van Bergen for practical assistance during the experiments; Marleen van Eijk and Merijn de Bakker for help with RNA isolation; Bart Pannebakker for helpful discussions on data analysis; Henri van der Geest for bioinformatic asssistance; the UCL Legion High Performance Computing Facility (Legion@UCL) and associated support services for computational support; Clara Lacy for the butterfly drawings; and Judith Mank and Stephen Montgomery for helpful comments and suggestions on the manuscript.

---

**Figure S1.**
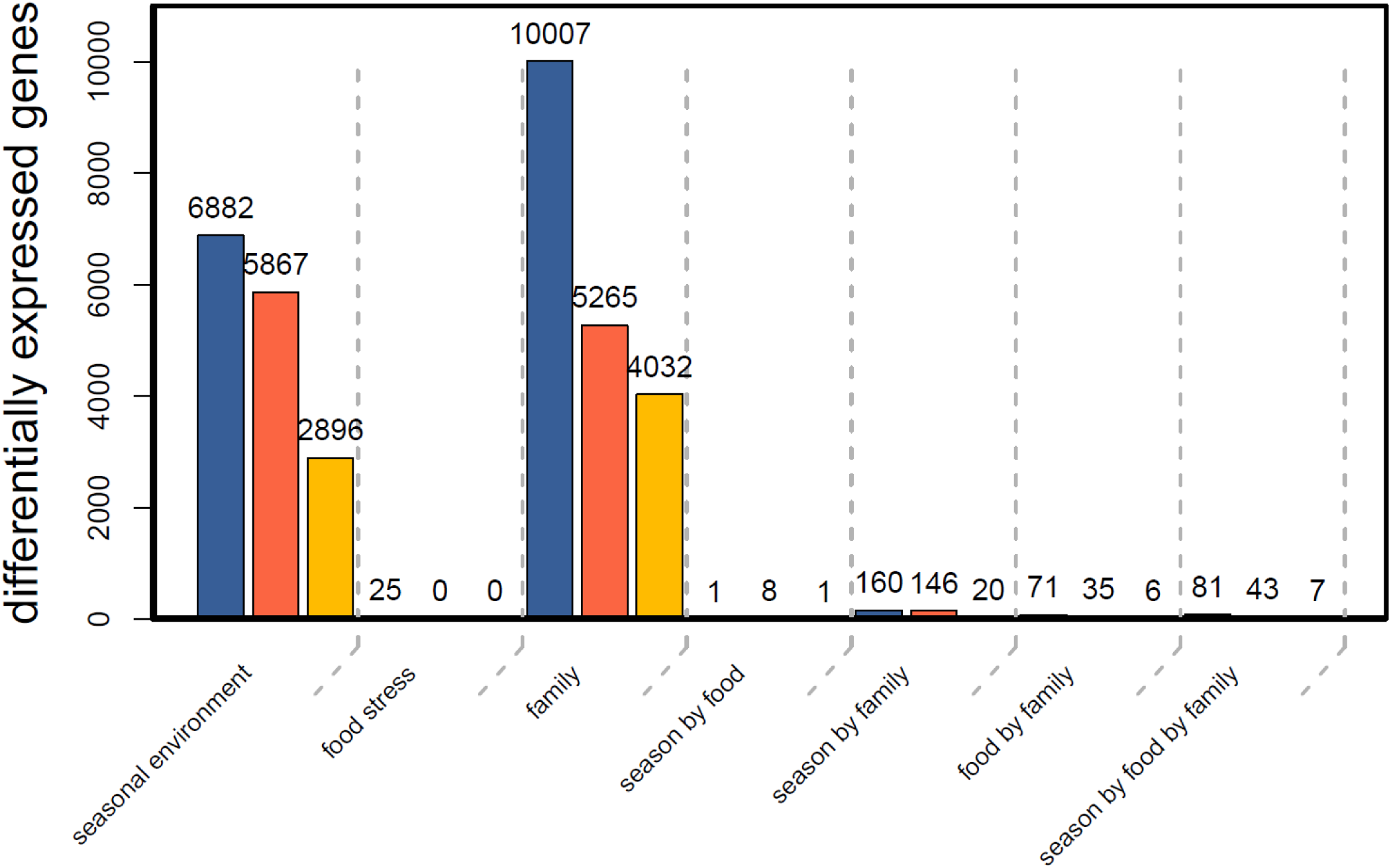
Suppl Fig S1.

**Figure S2.**
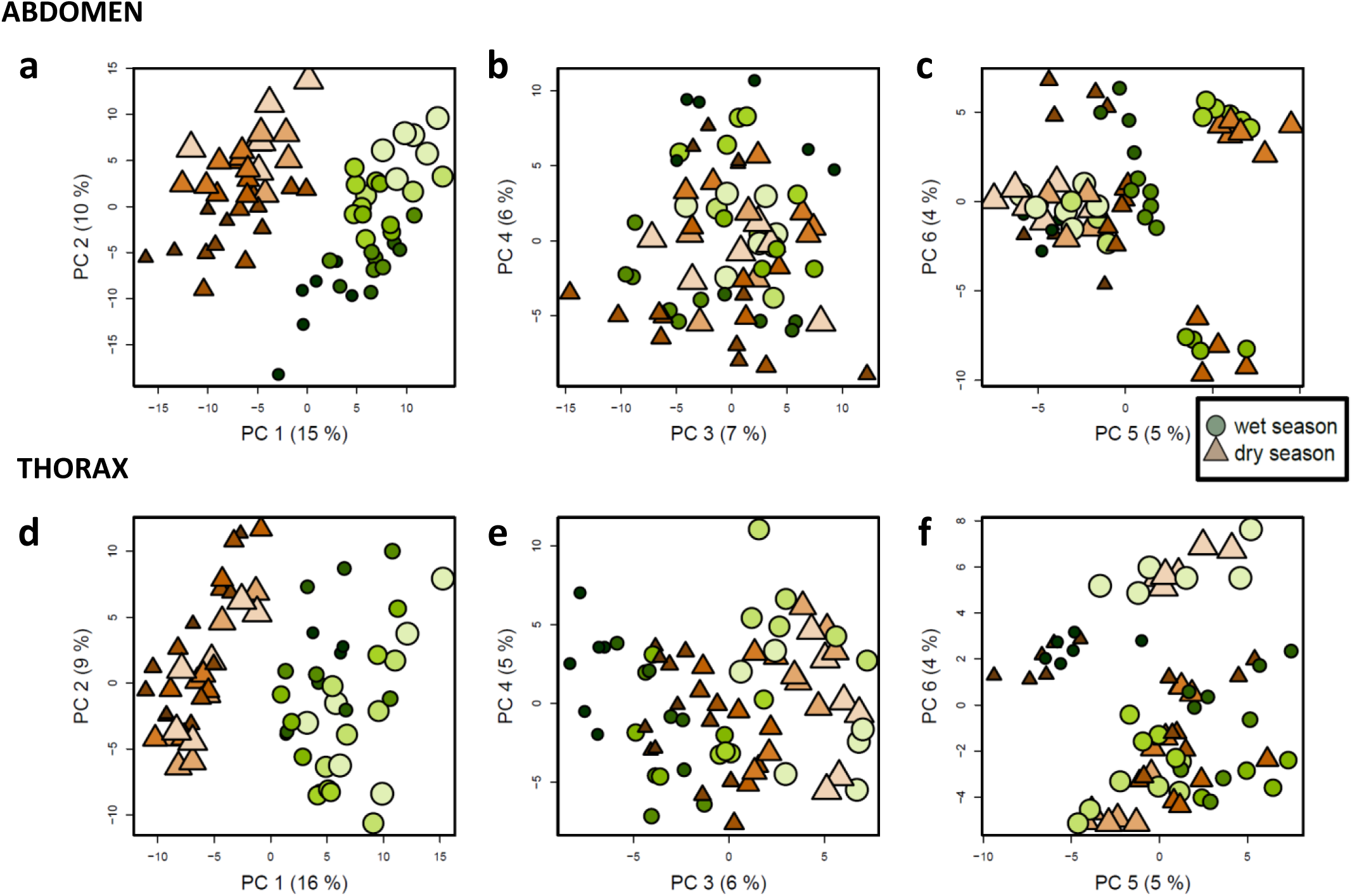
Suppl Fig S2.

**Figure S3.**
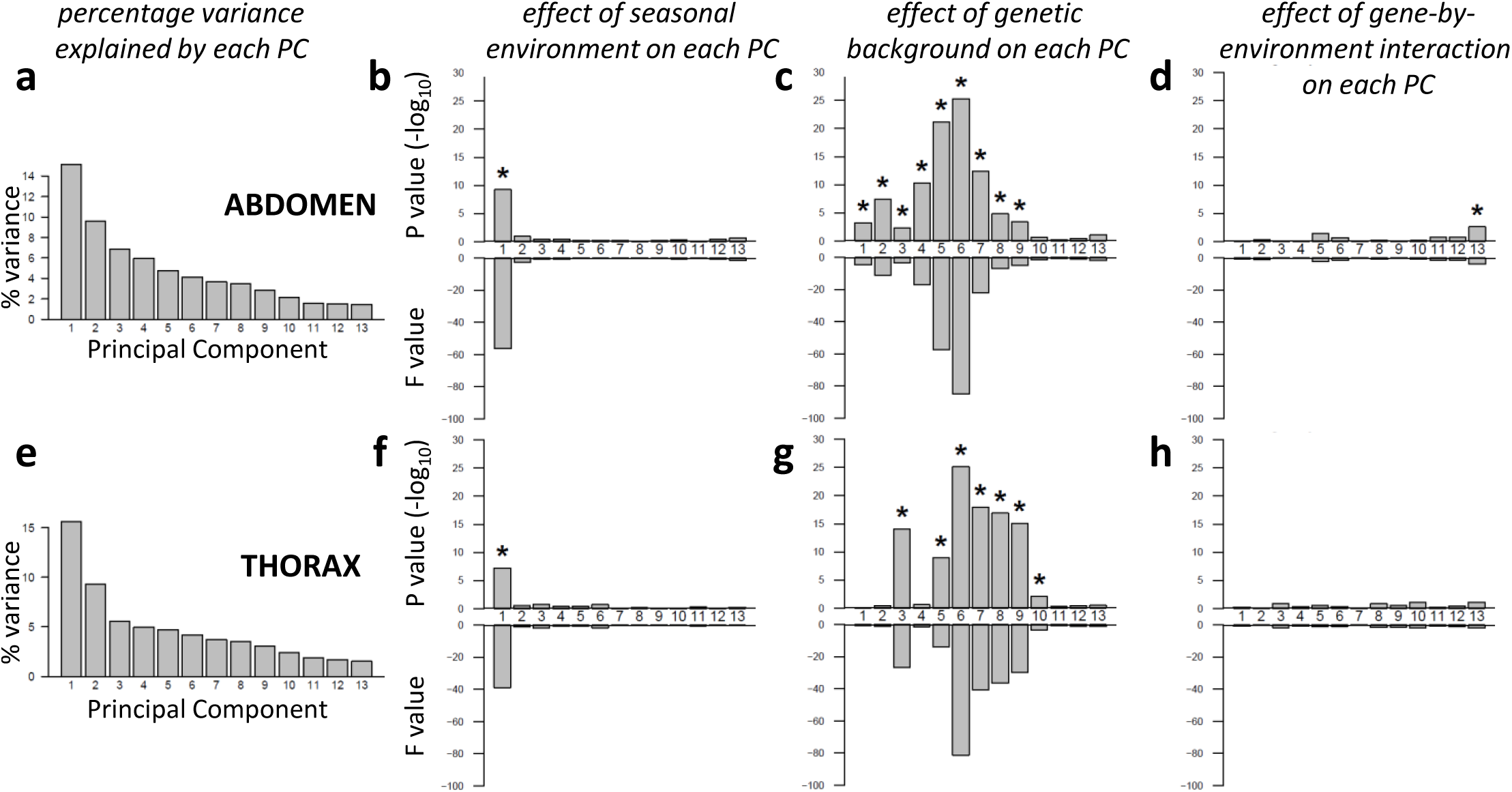
Suppl Fig S3.

**Figure S4.**
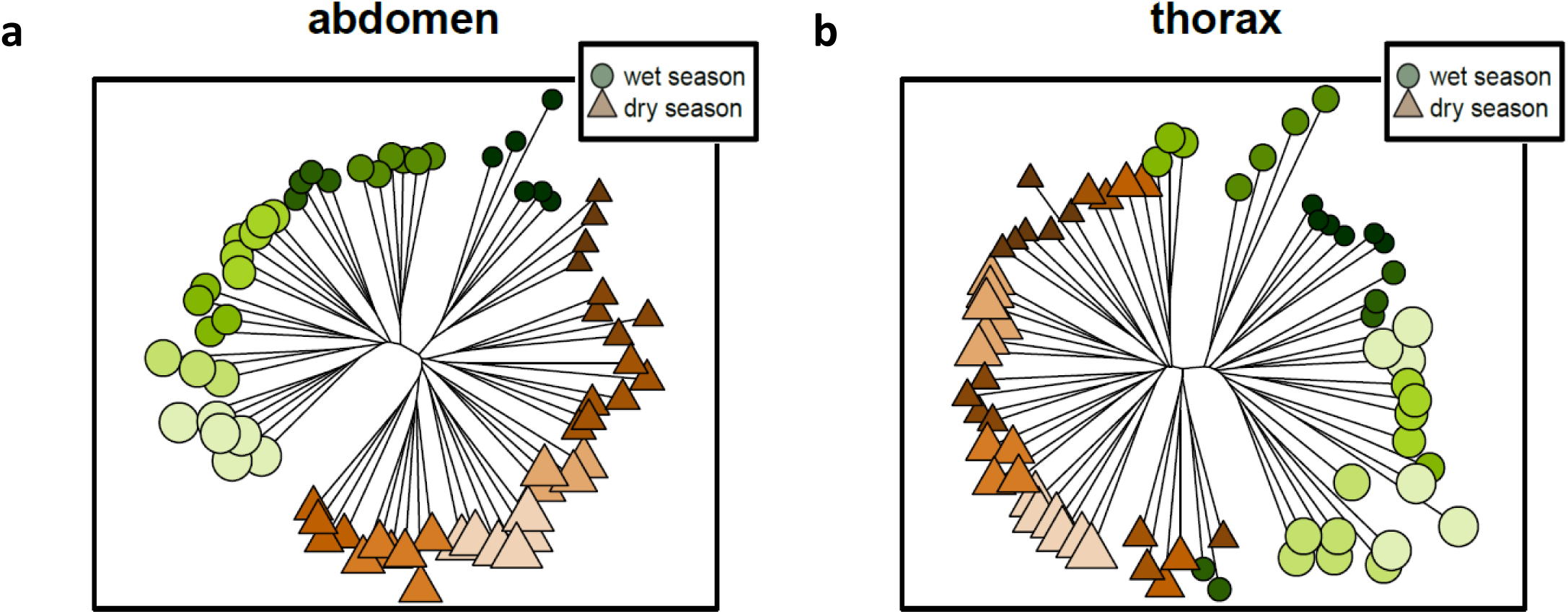
Suppl Fig S4.

**Figure S5.**
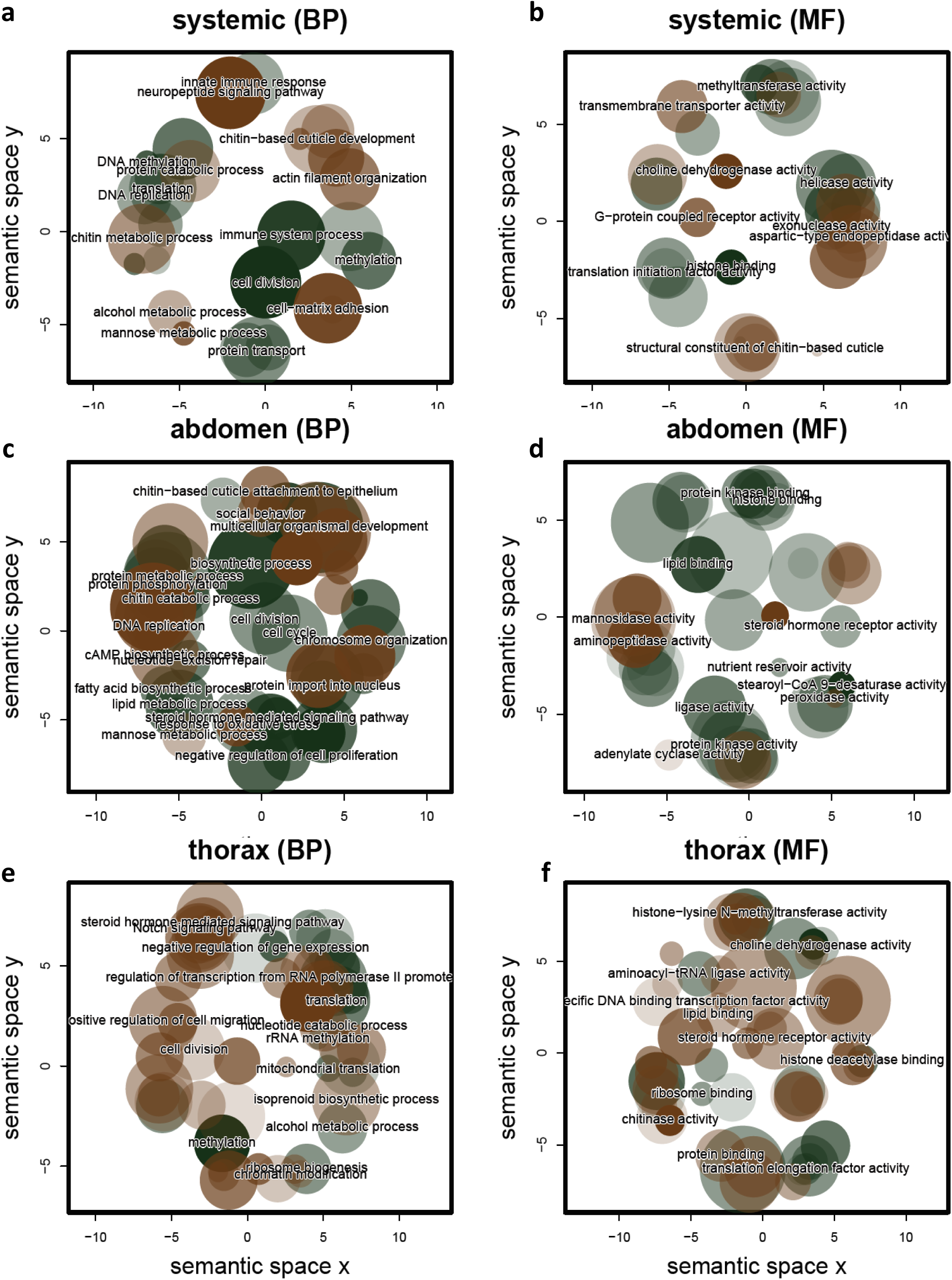
Suppl Fig S5.

**Figure S6.**
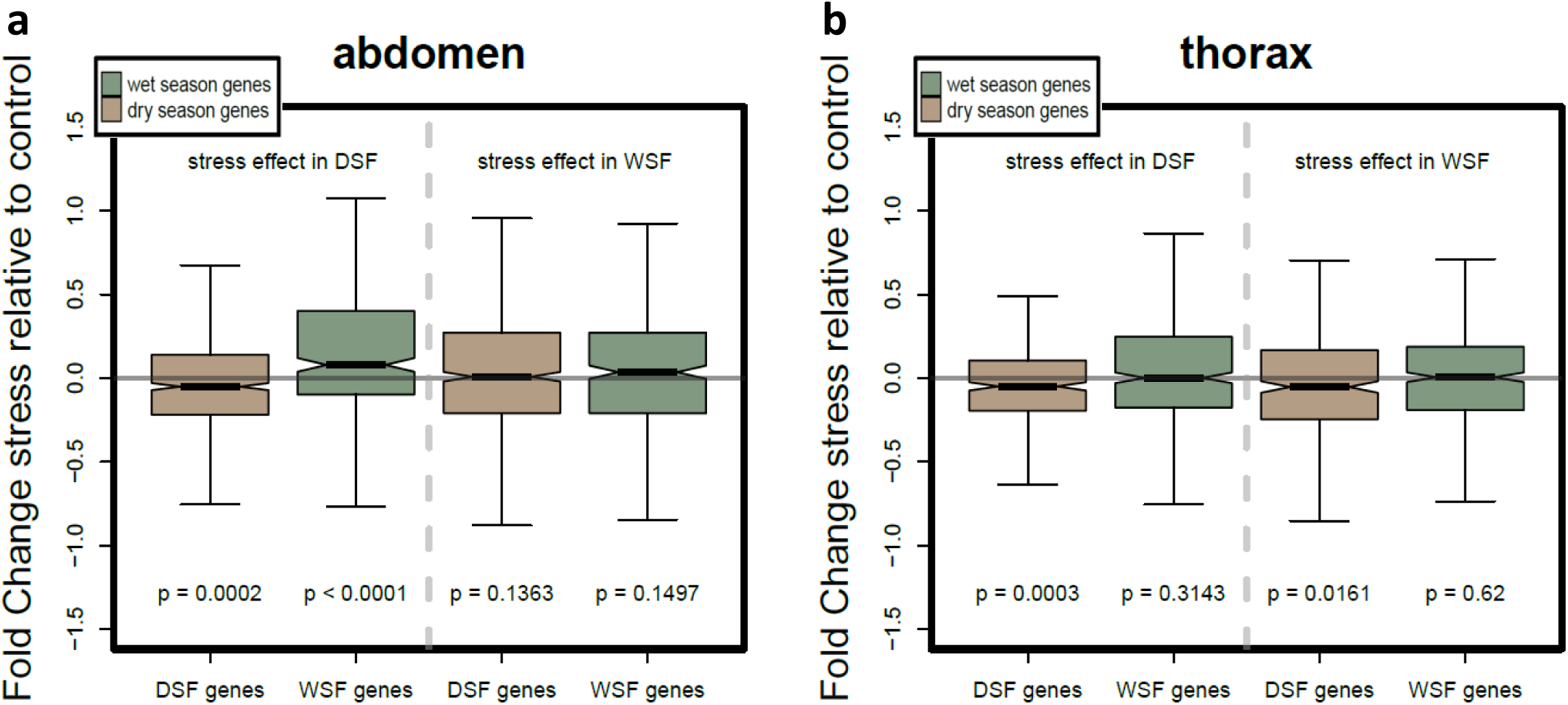
Suppl Fig S6.

**Figure S7.**
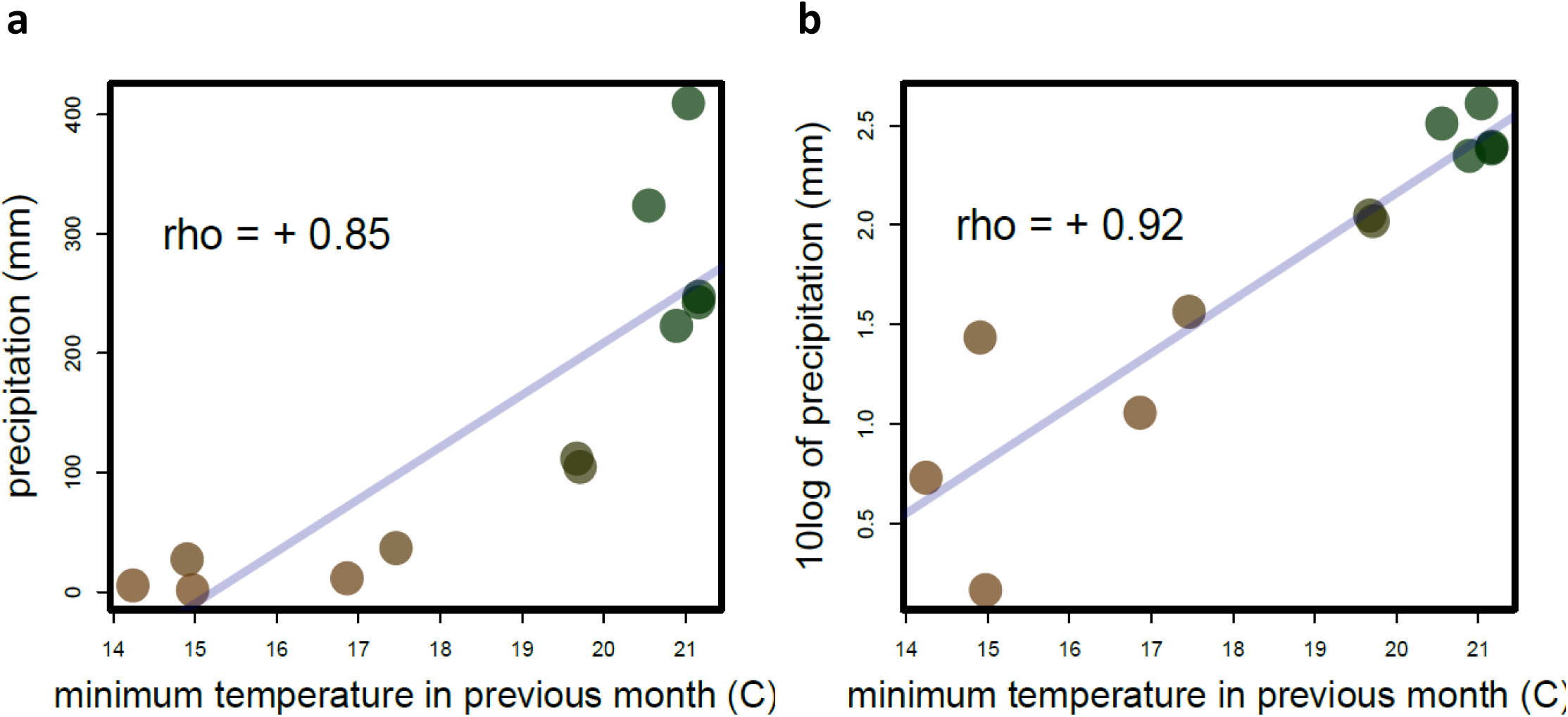
Suppl Fig S7.

